# Biotic interactions explain seasonal dynamics of the alpine soil microbiome

**DOI:** 10.1101/2023.04.17.537150

**Authors:** Anna Maria Fiore-Donno, Jule Freudenthal, Mathilde Borg Dahl, Christian Rixen, Tim Urich, Michael Bonkowski

**Affiliations:** Terrestrial ecology group, University of Cologne, Cologne, Germany; Institute of Microbiology, University of Greifswald, Greifswald, Germany; WSL Institute for Snow and Avalanche Research SLF, Davos Dorf, Switzerland; Climate Change, Extremes and Natural Hazards in Alpine Regions Research Centre CERC, Davos Dorf, Switzerland

**Keywords:** biotic interactions, alpine ecosystems, protists, metatranscriptomics, soil food web

## Abstract

**Background:** The soil alpine microbiome is dependent on season and elevation, yet there is limited understanding of how complex communities are differentially shaped by abiotic and biotic factors. Here we investigated the spring-to-summer dynamics of soil microbiomes in alpine grasslands, focussing on soil food web interactions. To this end, we conducted a survey along altitudinal transects in three mountains in the Alps, in spring at snowmelt and in the following summer, recorded vegetation and topographic, climatic and edaphic parameters for 158 soil samples. By using metatranscriptomics, we simultaneously assessed prokaryotic and eukaryotic communities, further classified by nutrition guilds.

**Results:** Our results show: (i) that biotic interactions could explain more variation of the microbial communities than topographic and edaphic variables, more for consumers than for preys, and this effect was stronger in summer than in spring; (ii) a seasonal dynamic in biotic interactions: the consumers’ pressure on preys increases from spring to summer, resulting in a higher diversity and evenness of preys.

**Conclusion:** In alpine grasslands, consumers effectively contribute to maintain the diverse soil bacterial and fungal community essential for ecosystem functioning.

## 1 Background

Understanding seasonal dynamics of the soil microbiome is an important step towards modelling the effect of climate warming, i.e. if and how the balance between the carbon stored in soil and the CO_2_ released to the atmosphere will be altered [1]. It is not yet clearly understood how the main components of the soil food web - bacterial and fungal primary decomposers and their main consumers (predatory bacteria, heterotrophic protists and bacterivorous and fungivorous nematodes) - are structured in alpine regions, and in particular which of the abiotic drivers, elevation or season, has a preponderant influence [2, 3]. Along altitudinal gradients, bacterial and fungal diversity generally decreases with elevation [4], while protistan diversity increases [3]. It is becoming gradually clear that in considering only high-rank taxonomic groups (i.e. bacteria or fungi), changes in community composition may be overlooked, since trophic guild’s opposite responses may cancel each other out [5]. Indeed, specific taxa or functional groups of bacteria, fungi and protists have different optima along environmental gradients and therefore are differentially affected by seasonal and/or altitudinal changes [4, 6-8]. For example, bacteria and fungi can show a diversity optimum at high (e.g. Verrucomicrobia, Beta- and Deltaproteobacteria)or middle altitude (Chloroflexi and most fungi) [7]. Similarly, distinct factors drive protistan nutritional guilds [9], e.g. consumers, parasites and phototrophs differentially respond to altitude and edaphic factors; noteworthy, only 40% of the species turnover of these communities could be explained by abiotic factors [8].

While most studies agree on the importance of pH and elevation for shaping bacterial communities in general, the unexplained variance is usually larger for fungi and protists [3, 4, 7]. Recently, it has been shown that models integrating biotic interactions in addition to abiotic environmental variables explained significantly more variation in microbial metacommunity assembly [10-12]. Biotic interactions are composed of two main types, competition for resources (mostly occurring among species at the same trophic level) and predation, both potentially contributing to promote species co-existence or exclusion [13]. Thus, including biotic interactions is of great importance for process-based understanding and forecasting ecological responses [14].

Our study aimed at disentangling the interactions of seasonal, topographic, edaphic and biotic factors and how they shape the soil microbiome. To this end, we used metatranscriptomics to simultaneously assess the prokaryotic and eukaryotic microbial diversity in alpine grasslands in Switzerland, near Davos (Fig 1a & b), at a regional scale (158 samples on three different mountains), taking into account elevation (altitudinal transects from c. 1900 to 2800 m above sea level) and season (spring and summer; Fig. 1c,d & e). The small subunit ribosomal RNAs (16S and 18S), assigned to the genus level, were used to identify the bacterial, archaeal, fungal, metazoan and protistan communities. To investigate biotic interactions, identified genera were further classified, when possible, according to nutrition type and lifestyle.

**Fig. 1.**
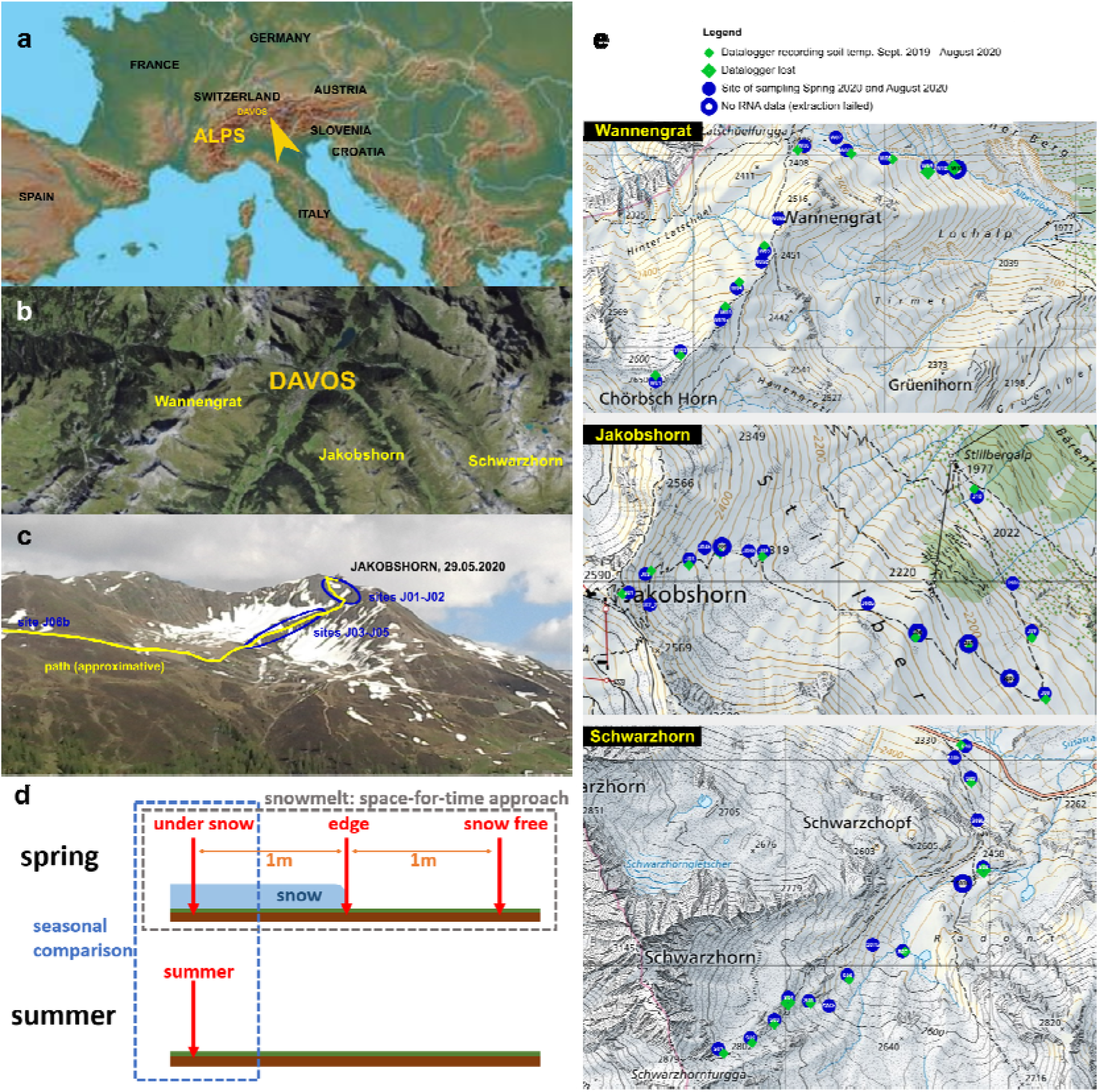
| Maps of the collection sites and sampling scheme. **a**, The Alps and Davos in Europe. **b**, The three mountains where the study took place. **c**, View of the sites at the top of the Jakobshorn during the spring sampling. **d**, A scheme of the sampling design per site. **e**, Sites and dataloggers position along the altitudinal gradient in each mountain.

We addressed the following major questions: (i) which are the dominant components of the soil microbiome during snowmelt and what is the fate of these communities in summer? (ii) do consumers contribute to the seasonal turnover of the bacterial and fungal communities? (iii) which are the main drivers of the soil alpine communities, topography, soil properties, seasonality or biotic interactions? We hypothesized that consumers would have a major role in shaping preys’ communities, with an increase of consumers and a decrease of preys during the season, and that season would play a more important role than elevation. Finally, we hypothesized that each nutritional guild would differ in its response to environmental and seasonal changes.

## 2 Materials and Methods

### 2.1 Sampling design

The study area is located around the city of Davos in the eastern central Alps in Switzerland (Fig. 1). The climate in this region is characterized by a mean annual temperature of 3.5° C, 193 days of frost, a total annual precipitation of 1,022 mm and fresh snow on 69.1 days a year [15]. The mineral bedrock is composed of acidic rocks, mostly gneiss and mica schist [16].

Sampling was designed to capture the biogeochemical changes happening at snowmelt, in alpine grasslands at altitudes comprised between 1,972 and 2,816 m. For this, in spring 2020, we selected 15 - 16 sites per mountain, where patches of snow were still present. In each site, we collected three soil samples constituting a time-series: the first sample was still under the snow, the second at the edge of the patch, thus during snow melt, and the third in soil recently freed from snow (Fig. 1d). The three samples were one meter apart from each other. To assess a medium-scale reproducibility of our data, we repeated the same scheme in three mountains: Jakobshorn, Wannnengrat and Schwarzhorn, while the altitudinal gradient would allow comparing the biogeochemical data between early and later snow melting times. To establish a seasonal cycle, we have conducted a further sampling on the very same sites that were under the snow in spring in August 2020.

### 2.2 One-year soil temperature records

In each mountain, ten dataloggers (ibutton DS1922L, Maxim Integrated Products, San Jose, CA, US) set up to record the temperature every 7,200 s with a precision of 0.0625 °C were placed early September 2019 c. every 50 m along the altitudinal gradient. They were inserted into an inflatable balloon for protection, attached to a 10 cm screw and placed at c. 5-8 cm into the soil, with a visible aluminium label. They were recovered during the summer sampling, end of August 2020. (Supplementary Fig. 1). Coordinates and altitudes were estimated with a GPS (Trimble Geo XH 6000, Trimble Inc. Sunnyvale, CA, US) with a precision of few cm (Supplementary Table 1).

### 2.3 Collection of soil samples

The spring collection took place between the 19.5 to the 24.6.2020, the three highest sites in the Schwarzhorn could only be collected the 6^th^ of July. Summer sampling took place in the following August, from the 20^st^ to the 25^th^. Soil temperature at the moment of collection was recorded using the same dataloggers as above, inserted at a depth of c. 5-8 cm, and left for c. 10 min. to record the temperature with a precision of 0.0625 °C (Supplementary Table 1). Two to 4 g of wet soil for RNA extraction were collected with a clean plastic spoon and immediately put in a sterile, RNAse-free 15 ml plastic tube containing 6.5 ml of Life Guard soil RNA preservation (Qiagen GmbH, Hilden, Germany) and stored in an insulated box with cooling packs. The samples when later stored at -20 °C as soon as we were back at the WSL Institute in Davos. They were not allowed to thaw until extraction at the University of Greifswald or Cologne. Soil (c. 200 g) for determining edaphic properties was collected using a clean metal spoon and stored at 4 °C in a polyethylene bag.

### 2.4 RNA extraction, reverse-transcription, library preparation and sequencing

Prior starting the following steps, great care was taken to work in an RNAse-free environment, notably by treating all objects that would come in contact with the samples with RNaseZap, RNase decontamination solution (Sigma-Aldrich, MO, USA). The tubes containing the soil samples in Life Guard were defrosted, centrifuged and the buffer removed. Circa 1 g of wet soil was removed with an autoclaved and RNAse-free treated spatula to be put in the tubes of the RNeasy PowerSoil Total RNA kit (Qiagen GmbH, Hilden, Germany). The manufacturers’ protocol was strictly followed, apart for the disruption step, which was carried on a MP Biomedicals FastPrep-24 homogenizer for 30 s at 5 m/sec. The RNA was eluted in 50 μl of the buffer SR7, with the addition of 1 μl of recombinant RNasin ribonuclease inhibitor (Promega, Madison, WI, USA). Quantity and quality was roughly evaluated on an agarose gel. DNAs were digested with DNAse I (New England BioLabs, MA, USA) following the manufacturers’ protocol. DNase was not inactivated, since we proceeded immediately to remove proteins and small RNAs using the Megaclear kit (Invitrogen, CA, USA), following the manufacturers’ protocol. Samples were eluted with 50 μl of preheated elution buffer, and quantified using a Qubit 30 Fluorometer (Invitrogen, CA, USA) using 2 μl of the RNA in the high sensitivity buffer. Quality was estimated for some samples with a 2100 Bioanalyzer (Agilent, CA, USA) using the Prokaryote Total RNA Nano assay. Samples with a concentration < 11 ng/μl were precipitated with 1:10 volume of 5M Ammonium Acetate and washed with ethanol, following the protocol of the Megaclear kit. Samples with a RNA concentration >10ng/µL were further processed. Libraries were prepared using the NEBNext Ultra II Directional RNA Library Prep Kit (New England Biolabs, Ipswich, MS, USA) with no removal of rRNAs or selection of mRNAs. The first strand cDNA synthesis incubation time at 42°C was increased from 15 to 50 min. To select fragments of cDNA (after the second strand synthesis) of 370-600 bp, the fragmentation time was reduced to 10 min. The library size selection option "400bp" was chosen and the final libraries were amplified with 12 PCR cycles. Libraries were sequenced with a single complete run of NovaSeq SP FC (Illumina Inc., San Diego, CA, US), length of paired sequences of 250 bp at the Cologne Genomic Centre, University of Cologne, Germany.

### 2.5 Sequence analyses - filtering and identification

From the Illumina NovaSeq sequencing 1.02 x 10^9^ raw sequences were obtained. These were submitted to the PhyloFLASH processing pipeline following default settings [17]. In brief, the pipeline identifies SSU rRNA sequences by aligning unpaired sequences to a filtered SILVA database (v.138, NR99), from which LSU, low-complexity and cloning vectors fragments have been removed. SSU sequences were identified with a short sequence aligner for DNA and RNA-seq data (BBmap; https://sourceforge.net/projects/bbmap/, last accessed Mai 2022), with the default setting of a minimum identity of 70%. For Bacteria and Archaea, taxonomic affiliation was assigned by taking the last common ancestor (LCA) of the taxonomy strings of all the database hits, using the SILVA taxonomy. Eukaryotic forward and reverse SSU sequences were assembled using FLASH [18] and low-quality sequences were filtered out with default settings. Eukaryotic sequences were identified to the genus level using a slightly modified PR2 database [19], using Blast + [20] with a e-value of 1e-10 and keeping only the best hit. Unicellular eukaryotes were classified as protists, and additional information was added about lifestyle (free-living, plant or animal parasite) and nutrition (heterotroph, autotroph or myxotroph), whenever possible, according mostly to Adl et al. (2012). Nematodes were classified as eukaryvore, bacterivore, plant-feeding, fungivore, animal parasite or omnivore according to [21] and http://nemaplex.ucdavis.edu/IndexFiles/etol.html (last accessed June 2022). Fungi were classified as saprotroph, lichen, lichen parasite, plant parasite, animal parasite, fungal parasite, ectomycorrhizal, endomycorrhizal, ericoid mycorrhizal, endophyte, symbiotroph and animal endosymbiont according to [22] (Supplementary Table 2). To assess the biotic interactions, we considered broadly-defined groups of consumers and preys. In the consumers were included the bacterial predators (phyla Myxococcota and Bdellovibrionota, 94.7% of the assemblage), the heterotrophs and free-living protists (4.3%), and the free-living nematodes (1%). The preys were composed of all other bacteria (99.2%), fungi (0.77%) and the autotrophic protists (unicellular algae, 0.03%).

### 2.6 Edaphic properties and vegetation survey

Soil water content, pH, Soil organic C and total N, soil microbial biomass C and N, dissolved organic C and total dissolved N were measured as already described [23] (Supplementary Table 1). Vegetation was recorded from June to August in 2020 and 2021, inside a circle of 40 cm diameter around the spot where the soil sample was taken. All vascular plants rooted inside the surface were identified according to [24], and the percentage of the surface they covered was estimated (Supplementary Table 1).

### 2.7 Statistical analyses

All statistical analyses were carried out within the R environment (R v. 4.1.3) [25] on the taxonomic abundance/samples (Supplementary Table 2) and on the samples topographic, edaphic and biotic characteristics (Supplementary Table 1). Unless otherwise specified, community analyses were performed with the package vegan 2.5-7 [26]. **Alpha- and Beta-diversity**: To evaluate if more sampling and sequencing effort would have revealed more taxonomic richness, we carried out an analysis based on accumulation curves (function *specaccum*) and rarefaction curves (package and function iNEXT 3.0.0), using the abundance table, with a 97% confidence interval, 50 bootstraps and 50 knots; the latter function also calculates species richness (observed and estimated) and sample coverage. Exponential Shannon indices were calculated with the function *renyi* (with the hill parameter), so that they are more comparable between samples. Significant differences in sequence counts, alpha diversity and evenness between seasons were determined by analysis of variance and Tukey tests (package agricolae 1.3-5, function *aov* and *HSD.test*) with a p-value ≤ 0.05, while correlations between the same data were determined with the function *lm*.

Beta diversity between mountains, altitudes as factors and snow coverage in spring was inferred by Principal Coordinate Analysis (PCoA, function *cmdscale*), using Bray-Curtis dissimilarities (function *vegdist*, method “bray”) on the relative abundances of taxa of interest (Bacteria, Archaea, Protists, Fungi and the functional groups consumers and preys), then plotted with the package ggplot2 3.3.5. Principal component analysis revealed an influence of altitude and mountain on bacterial beta-diversity (and thus that of preys, since bacteria dominate by far this assemblage), but this effect was driven mostly by the three highest sites in the Schwarzhorn (Supplementary Fig. 2). With these outliers removed, a decrease of the variation explained by the first axis was observed (40.1 to 31.3%), and only a slight trend for altitude was visible. Fungal, metazoan, protistan and consumers’ communities did not display a clear trend, only the protistan summer communities and some high-altitude sites tended to be different (Supplementary Fig. 2). Variation partitioning (function *varpart* applied to the Hellinger-transformed taxa dataset and using RDA, function *rda*) was applied to assess the amount of explained beta diversity by the factors mountain, altitude and snow coverage in spring.

Differential abundances of the most abundant taxa across seasons were calculated with the package DESeq2 1.30.1 [27]; DESeq objects were created with function *DESeqDataSetFromMatrix* and normalized (function *estimateSizeFactors*), then the differential expression was calculated using *DESeq* with parameters minReplicatesForReplace = Inf, sfType = "poscounts". Results with an adjusted p-value <0.01 and an absolute log2fold >0.5 were considered as significant and plotted with ggplot2 (*geom_segment*).

Distance-based redundancy analysis (dbRDA, function *dbrda*) was conducted to describe the influence of environmental factors (scaled with function *scale*) on the distribution of the abovementioned taxa of interest (normalized by relative values by samples, function *decostand*). The most influential variables were identified with the function *ordistep* based on the Akaike Information Criterion (AIC), and the resulting model tested with *anova*.

To estimate the proportion of variance of topographic (mountain and altitude), biotic (Shannon vascular plants, Shannon preys or consumers) and edaphic (soil temperature, water content, pH and organic C) factors on the diversity of consumers and preys by season, variance partitioning analyses were conducted. The variance attributed to each category of factors (topographic, biotic and edaphic) and their intersections was estimated using the R^2^ of the linear models (function *lm*) of log-transformed Shannon indices of preys or consumers versus the other categories of factors.

To investigate associations between consumers and preys and inside consumers, networks analyses were performed on the taxonomic and functional table (Supplementary Table 2), first selecting the most abundant taxa (Supplementary Table 3) and then, for each season, applying a prevalence filter, i.e. selecting taxa that were present in more than one third of the sites. The filtered-out taxa were binned into a "pseudo-taxon", used for inferences for not altering the ratio between taxa, but not shown in the final results. The co-occurrence network was calculated at the genus level using FlashWeave [28], a package implemented in julia v.1.7.1 [29] with parameters for homogeneous data (sensitive=true and heterogeneous=false). The network was then summarized (using R) to higher taxa and functions and visualized using Cytoscape v.3.9.1 [30]. We excluded self-loops and associations inside preys.

## 3 Results

### 3.1 Metatranscriptomics

From the 158 collected soil samples, we obtained more than one billion RNA sequences, resulting in c. 8,000 identified taxa at the genus level, on average 2,349 per sample (Supplementary Table 4). Rarefaction curves describing the observed number of genera as a function of the number of sequences (Supplementary Fig. 3) suggested that our sequencing effort was sufficient, confirmed by the estimated sample coverage (between 99.95 and 100%, Supplementary Table 2). Accumulation curves describing the number of genera as a function of the number of samples, binned by altitude, mountain and snow coverage (Supplementary Fig. 4) did not reach a plateau and showed slight differences between environments, thus indicating differences in beta-diversity.

### 3.2 Taxonomic and functional diversity

Prokaryotes dominated by far the total assemblage, with 98.02% of the identified taxa, and among them Archaea were insignificant (0.06%). (Supplementary Table 2 and Supplementary Note 1). In bacteria, it is noteworthy that the predatory Myxococcota represented nearly 6% of the bacterial genera (Fig. 2a and Supplementary Note 1), thus the main consumer in soil, surpassing in abundance the heterotrophic protists. Among eukaryotes, Fungi and Metazoa accounted for 36% of the SSU rRNAs each, protists for 17%, and multicellular plants for 11% (Supplementary Table 2). More than one third of the protistan sequences were assigned to the phylum Amoebozoa (35%; Fig. 2b), including the very elusive slime-moulds Myxogastria. The phylum Rhizaria (23%) was in majority composed of Cercozoa (20%). Our functional assignment according to nutrition suggested that 88% of the protists were heterotrophs, 8% autotrophs and 2% mixotrophs. According to lifestyle, the majority was free-living (90%), 7% were animal and 2% plant parasites (Supplementary Table 2).

**Fig. 2.**
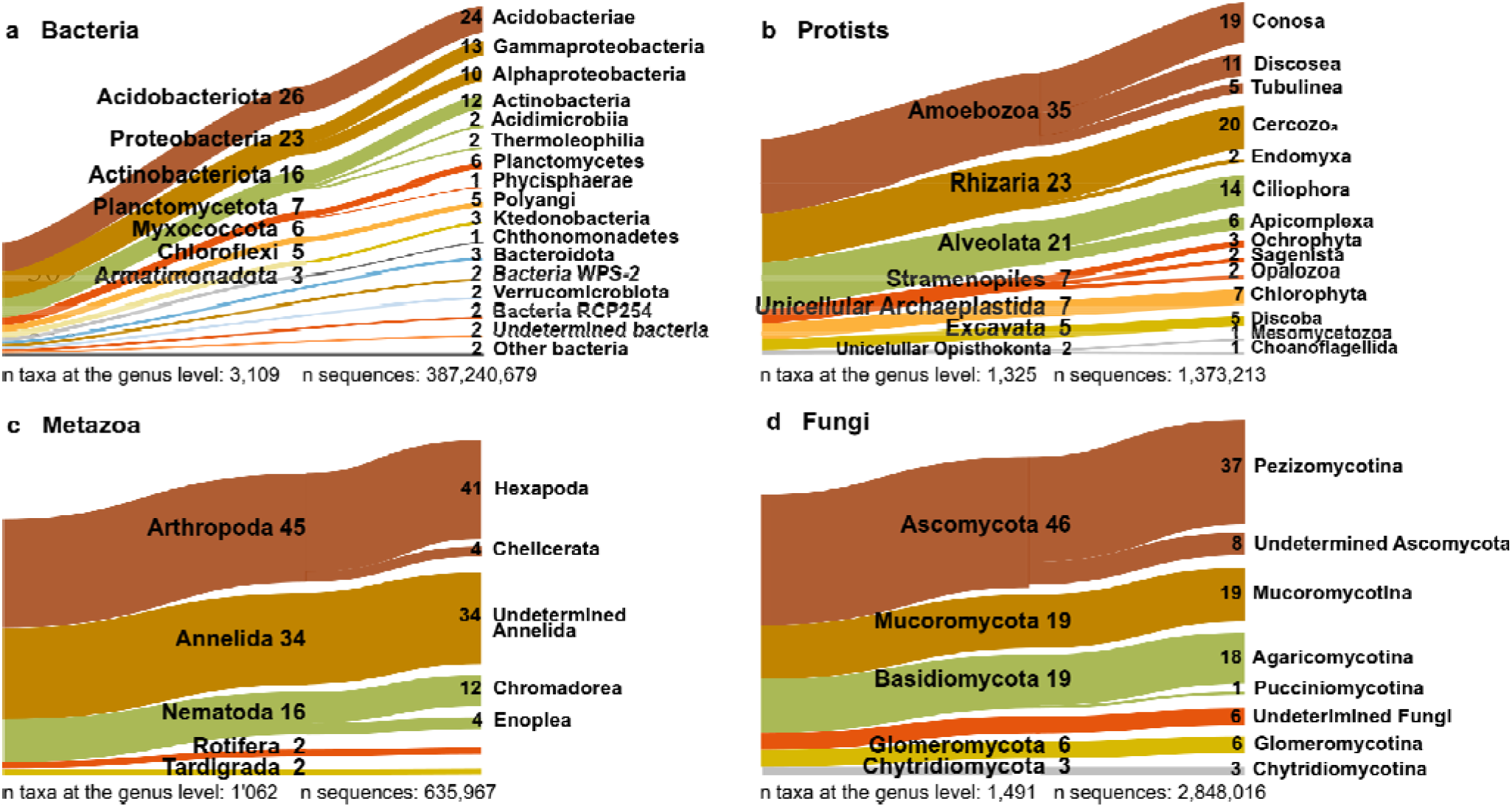
| Sankey diagrams showing the relative proportion of high-rank taxonomic groups, based on percentage of sequences identified to the genus level. Taxa < 1% are not shown. **a**, Bacteria; **b**, Protists; **c**, Metazoa; **d**, Fungi.

Among the animals (Metazoa), insects (41%) and ringed worms (34%) were most abundant (Fig. 2c), followed by nematodes (16%). Nematodes were classified as bacterivores (29%), plant-feeding (25%), fungivores (16%), eukaryvores (10%), omnivores (3%) and animal parasites (1%; Supplementary Table 2).

The fungi were dominated by Ascomycota (46%; Fig. 2d), of which mostly Pezizomycotina (37%), which also makes up most of Ascomycota in terms of described species. Of the 53% of the fungi that could be functionally identified, the largest group was saprotrophs (35%), followed by plant parasites (7%), endomycorrhizals (6%), ectomycorrhizals (5%), lichenized (5%), ericoid-mycorrhizals (2%), and other parasites and endosymbionts representing each <1% (Supplementary Table 2).

### 3.3 Soil temperatures and edaphic parameters

Soil temperatures (measured during the year in which the sampling took place) showed a day/night variation before and after the snow covered the soil; during winter under the snow cover, temperatures were stable and constantly stayed above 0 °C [23] (Supplementary Fig. 1). The snow cover lasted on average 228 days (max. = 286, min. = 186, sd = 21.2; Supplementary Table 1). Temperatures recorded during the sampling increased during snow melt, from under the snow (mean = 0.4 °C, sd = 0.2), to the snow edge (mean = 2.7, sd = 2.2) and the snow-free sample (mean = 10.2, sd = 4.2), and further increased in summer (mean = 14.1, sd = 4.7). Related to temperature and snow cover, soil water content decreased from under the snow (mean = 53%, sd = 15.6) to edge (mean = 49%, sd = 13.5) and snow-free samples (mean = 45%, sd = 12.9), and further decreased in summer (mean = 40%, sd = 12). The pH did not significantly vary during snowmelt. Microbial biomass and dissolved C increased during and immediately after snowmelt, as well as from spring to summer (Supplementary Table 1).

### 3.4 Dynamics during snowmelt and from spring to summer

In spring, despite of sharp environmental changes across the snowmelt gradient, no significant differences in the richness, diversity nor evenness of the communities of consumers and preys were observed (Supplementary Fig. 5). In contrast, richness, diversity and evenness varied between spring (the sample collected in spring under the snow) and summer (the sample collected at the same spot in August), but this varied differentially in pro- and eukaryotic taxa and functional guilds. For instance, the SSU rRNAs of consumers were significantly more abundant in summer than in spring, but no significant differences were found in diversity and evenness (Fig. 3a). In contrast, SSU rRNAs of preys were more abundant in spring, but reached their highest diversity and evenness in summer (Fig. 3b). These changes were mirrored when looking at the seasonal differences of the most abundant taxa: the number of sequences of Bacteria - mostly preys - decreased in summer, but the predatory Myxococcota increased (Fig. 3c). Fungi also decreased in summer, except the endomycorrhizal Glomeromycota (Fig. 3d). In Metazoa, all taxa were more abundant in summer, especially the arthropods (Fig. 3e). The increase of protistan abundance in summer was mainly due to Rhizaria, while Alveolata and Amoebozoa decreased in summer (Fig. 3f).

**Fig. 3.**
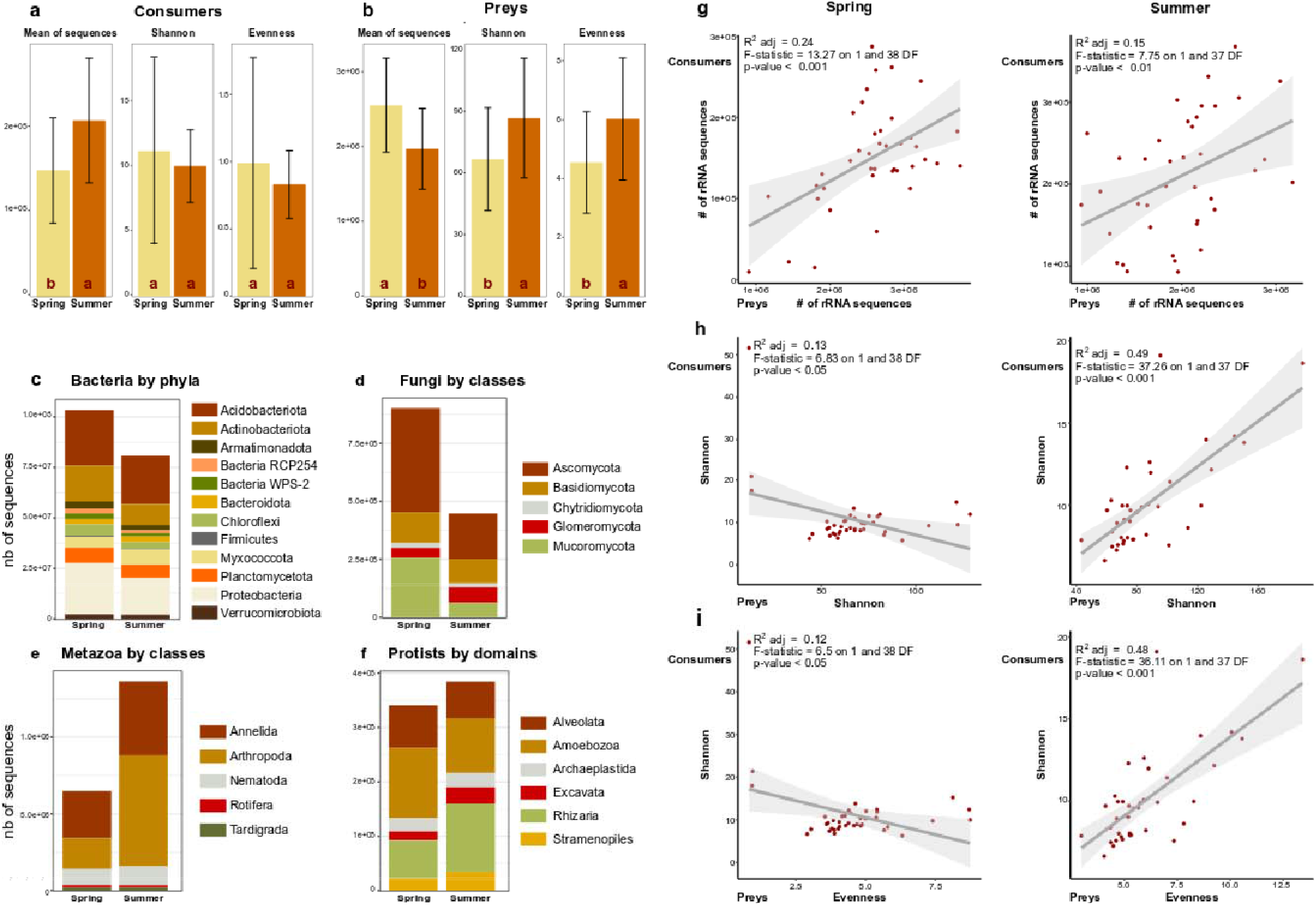
**| a-b**, Seasonal variation in the mean of SSU sequences, diversity (Shannon index) and evenness, from the sample under the snow in spring to summer. Significant changes (analysis of variance, p-value ≤ 0.05) are indicated by "a" (higher) and "b" (lower). Standard errors bars are shown. **a**, Consumers (predatory bacteria, heterotrophic and free-living protists, selected nematodes). **b**, Preys (non-predatory bacteria, fungi and autotrophic protists). Consumers increase from spring to summer (without significant changes in diversity or evenness); preys decrease, while their diversity and evenness increase. **c-f**, Seasonal differences in sequence counts. **c**, Bacteria, most abundant phyla. **d**, Fungi, most abundant classes. **e**, Metazoa, most abundant classes. **f**, Protists, most abundant domains. **g-i,** Linear correlations between consumers (y axis) and preys (x axis), in spring and summer. **g**, Number of rRNA sequences. **h**, Shannon indices. **i**, Shannon index of consumers versus evenness of preys. Dots = samples. Grey surface = 95% confidence interval.

Changes in abundance, diversity and evenness of consumers and their potential preys were correlated: there was a positive correlation in the abundance of consumers and preys, independently of the season (Fig. 3g). Despite of this, there were negative correlations between the diversity and the evenness of consumers with those of preys in spring (Fig. 3h-i). Contrastingly, the same negative correlations were strong and positive in summer (Fig. 3h-i). We questioned whether the increase in consumers in spring was related to the main abiotic changes occurring from spring to summer, i.e. a warmer and drier soil; we found positive linear correlations between soil temperature and water content and the richness of Myxococcota (F_[1, 156]_ = 7.6, p = 0.007) and Rhizaria (F_[1, 156]_ = 14.27, p < 0.001), but not for that of Amoebozoa.

### 3.5 Most influential environmental parameters

We tested how consumers and preys were responding to different factors, categorized as topographic (altitude, mountain), edaphic (water content, pH, soil temperature and organic C) and biotic (coverage and diversity of the vascular plants, diversity of Bacteria, Fungi, protists and consumers). Models (dbRDA) were estimated during snow melt in spring and from spring to summer (Supplementary Table 5). The models retained pH and the diversity of preys as the major drivers for consumers, in spring and summer; remarkably, the diversity of preys had more importance than pH in summer (Supplementary Table 5). Preys were also more influenced by the diversity of consumers (especially under the snow) than by topographic and edaphic factors. Bacteria were influenced by altitude and edaphic parameters in different combinations, depending on the samples; biotic factors had little influence. Less variation was explained in models estimated for protists, with low F-values and inconsistent results between samples. Fungi did not respond to topographic not edaphic factors, but sporadically to the diversity of bacteria and protists. No influence of the mountains was detected by this analysis (Supplementary Table 5).

For a reliable interpretation of our results, it was important to test the influence on the microbial communities of our sampling design, i.e. the four samples per site (Fig 1d), the altitudinal gradient (Fig. 1e) and the three mountains (Fig. 1b). The first axis of Principal component analysis only explained 31.3% of the variation of bacteria, with a slight effect of altitude. As in the dbRDA models, mountains had no effect, and fungal, protistan and consumers’ communities did not display a clear clustering trend with respect to the tested parameters (Supplementary Fig. 2).

### 3.6 Biotic interactions

Variation partitioning, conducted to disentangle the relative influence of topographic, edaphic and biotic factors, indicated a clear seasonal trend - all factors taken together explained more variance in summer than in spring, for both consumers and preys (Fig. 4a). A seasonal increase in the relative influence of biotic factors and biotic + edaphic factors on the variance of the communities was observed, higher for consumers (c. 17 times) than for preys (c. 2 times; Fig. 4b).

**Fig. 4.**
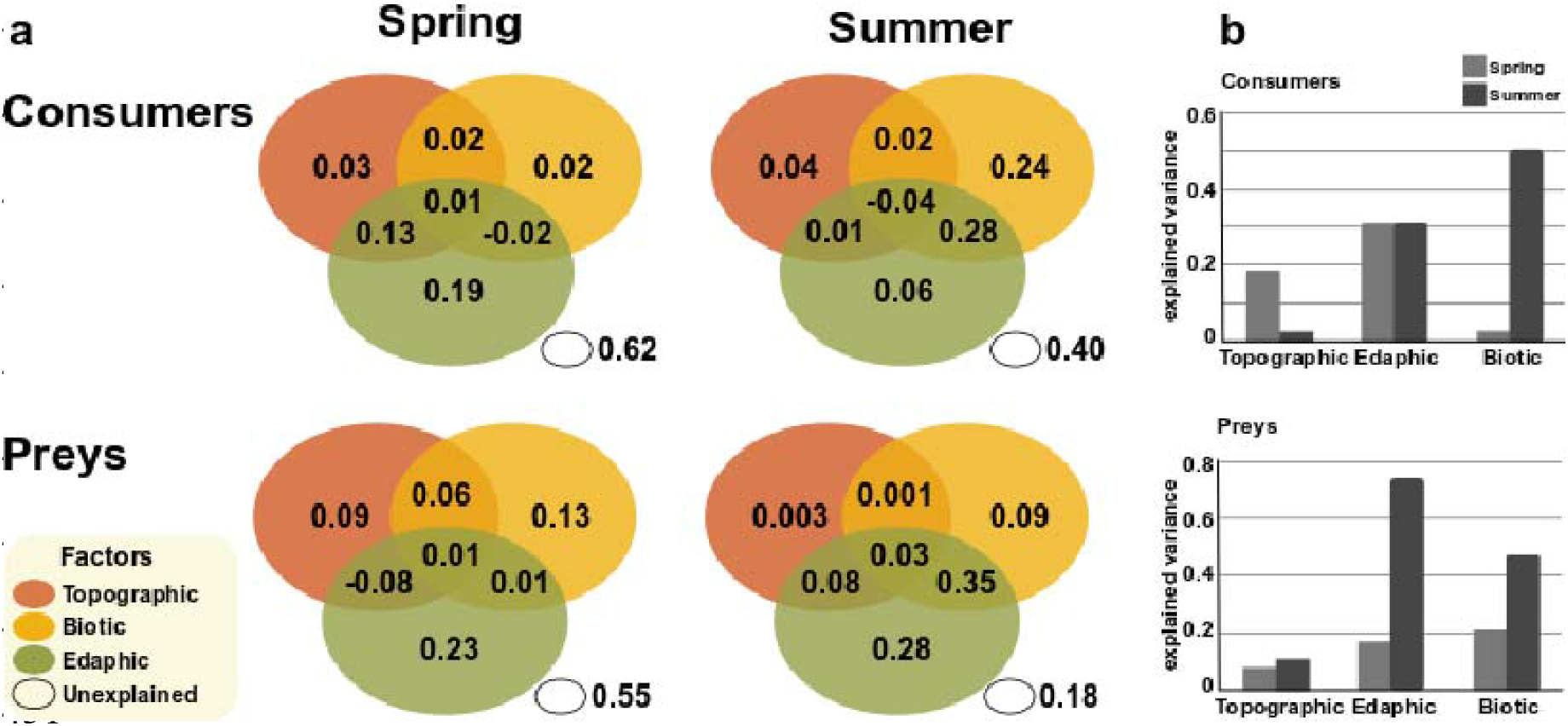
| Relative influence of topographic, biotic and edaphic factors on the diversity of consumers and preys, by season. **a,** The proportion of variance explained by topographic (upper left ellipse), biotic (upper right) and edaphic (bottom) factors, and the unexplained variance. **b,** Comparison of explained total variance by group of factors (the sum of each ellipse), in spring and summer.

Differential expression analysis revealed which taxa were significantly more abundant in spring or in summer. Strikingly, all groups selected by this analysis and more abundant in spring were preys (bacteria: Firmicutes, Actinobacteria and Proteobacteria, fungi: Ascomycota), while those more abundant in summer were consumers (bacteria: Myxococcota, protists: Rhizaria, Stramenopiles and Excavata; Fig. 5).

**Fig. 5.**
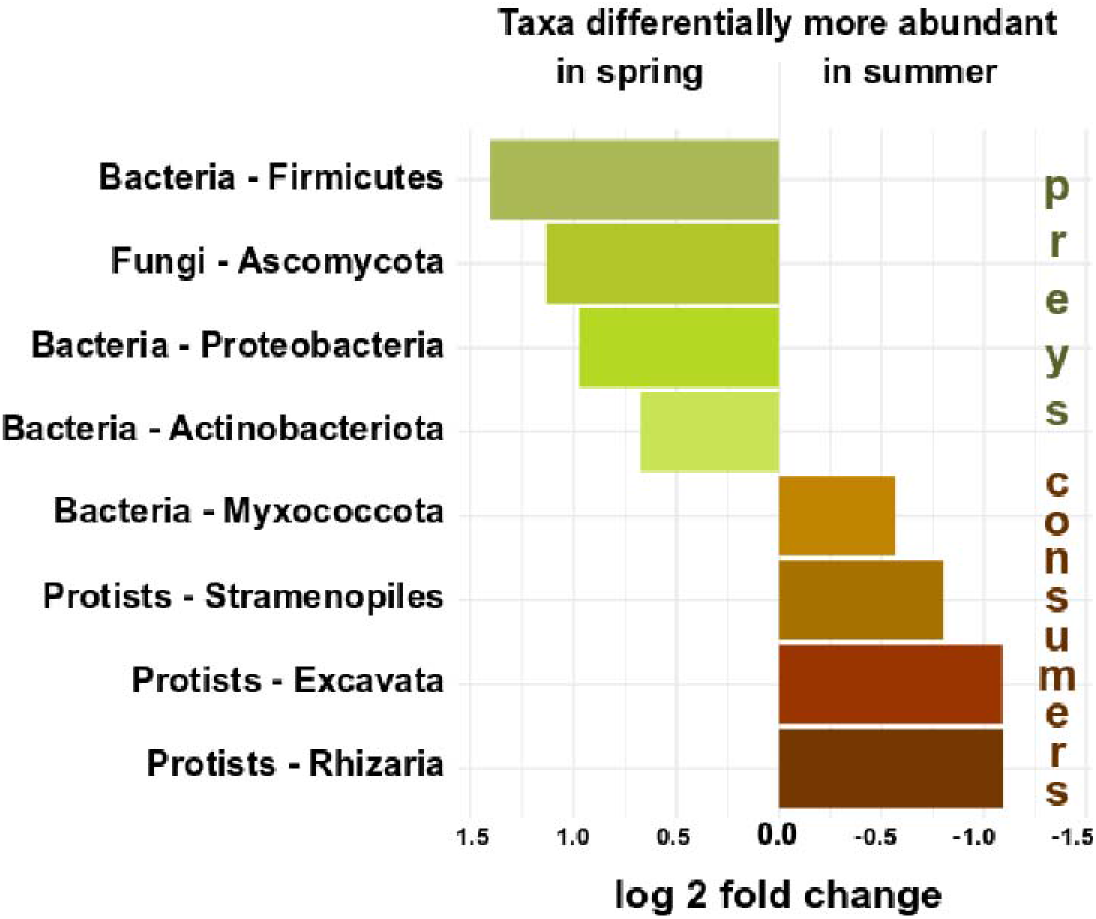
| Differential abundances. Taxa were filtered according to abundance and presence in one-third of the samples. All taxa differentially more abundant in spring and summer are preys and consumers, respectively.

The co-occurrence network confirmed the dominant role of biotic interactions between consumers and preys in summer, showing more associations - and more negative ones - in summer than in spring (Fig. 6a & b). In summer, Rhizaria stood out among the consumers with strong negative associations to several major bacterial phyla, i.e. Actinobacteria and Alpha- and Gammaproteobacteria. No seasonal differences were found in the networks of between-preys associations, which reflects competitive or facilitative interactions or the sharing of ecological niches (Supplementary Fig. 6). However, there are no noticeable differences in the structure of the networks, with changes mostly occurring in the proportion of the communities and the intensity of the associations.

**Fig. 6.**
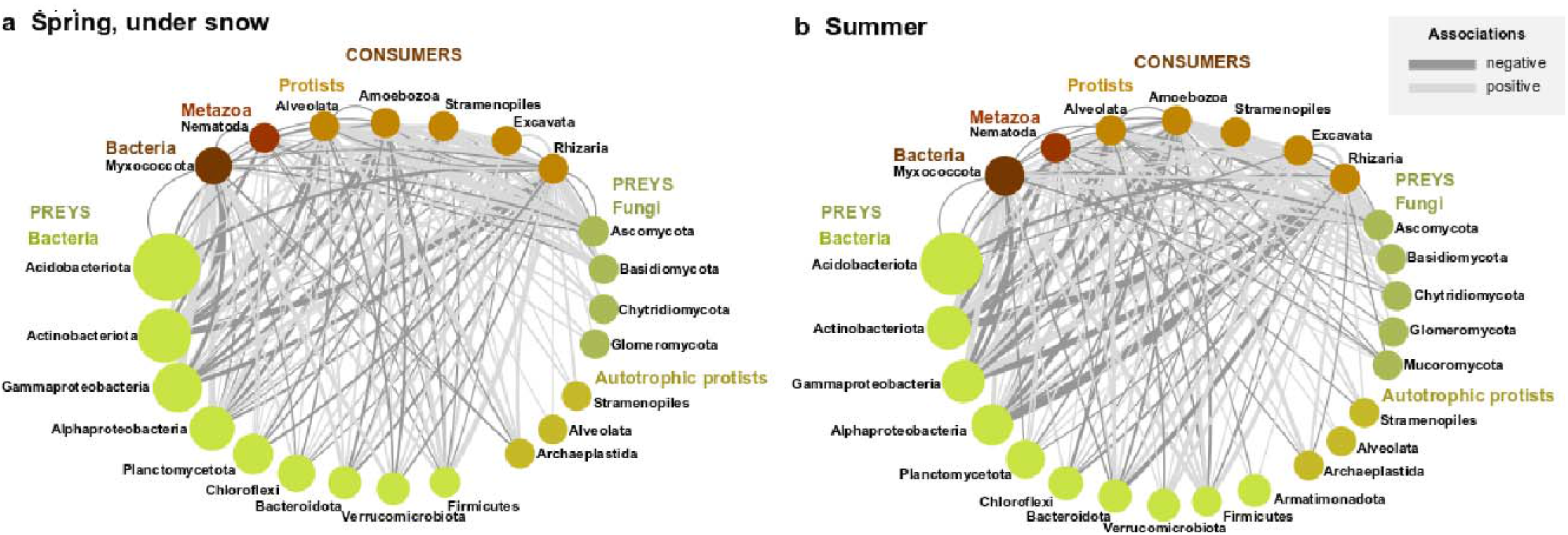
| Co-occurrence networks of abundant phyla of consumers and preys. **a**, Spring, under the snow. **b,** Summer. The size of the nodes (dots) are proportional to the number reads. Edges (connecting lines) represent positive (light grey) or negative (dark grey) correlations, with line width proportional to the number of correlations. Self-loops, taxa with a single edge and connections between preys are not shown.

## 4 Discussion

### 4.1 How biotic interactions shape the soil communities

The opposite and correlated changes in SSU rRNA abundance, diversity and evenness of consumers and preys from spring to summer strongly suggest an effect of predation: the increase of grazers’ abundance decreases the abundance of preys, while increasing their diversity and evenness - predators shape the preys’ communities (Figs. 3, 5 & 6). It has been repeatedly shown, especially in aquatic environments, that predation prevents competitive exclusion - i.e. the dominance of few better adapted species profiting from the resources of a given habitat [31-33]. Thus, as observed here, the increase in consumers’ abundance results in an increase in preys’ evenness and diversity, while without predators, competition for resources may lead to the dominance of fewer species [34]. The significance for ecosystem functioning is unmistakable: highly uneven communities, with an extreme dominance by few species, are less resistant to environmental stress [35]. Thus, both the number and relative abundances of species must be sustained to achieve a vigorous ecosystem functioning [36]. Here, we show that in alpine grasslands with a constant snow cover in winter, the spring to summer increase in consumers effectively contributes to maintain a diverse bacterial and fungal community.

The communities of consumers and preys were organised in highly inter-connected networks, with stronger negative interactions in summer (Fig. 6), which, added to the previous results, may likely reflect predator-prey interactions. Among protistan consumers Cercozoa and Amoebozoa display the strongest negative associations with major bacterial phyla. Supporting a consumer effect - but not proving it - the dominant cercozoan taxon is bacterivorous (Glissomonadida, 43% of all cercozoan sequences; Supplementary Table 2). Negative associations between Cercozoa and Actinobacteria (important polysaccharide decomposers) [37] and Alphaproteobacteria were also observed in the rhizosphere [38, 39], where the glissomonads were most abundant (compared to bulk soil and litter) [40]. A link between the decrease of Actinobacteria and the grazing of heterotrophic protists was also suggested [41].

However, negative associations could be purely abiotic, e.g. due to opposite sensitivity to environmental conditions, or biotic but driven by prey defences, e.g. Actinobacteria secreting secondary metabolites to avoid predation [42]. In addition to predation, competitive interactions have been shown to be a major driver in bacterial community composition [43], and fungal-bacterial competition explained 32% of the variance of planktonic bacterial communities [10]. Accordingly, we observed intricate networks between preys, not or only slightly affected by season (Fig. S6). However, we did not observe an increase of potential competitive interactions in bacteria with the increase of predation, as it was experimentally realized by the addition of bacterivorous nematodes [44].

Our results are in line with the numerous studies showing that bacterial communities were consistently driven by edaphic or topographic parameters (pH, altitude, water content and organic carbon), while eukaryotic communities showed weaker trends in response to environmental gradients compared to bacteria (Supplementary Fig. 2 & Supplementary Table 5) [4, 45]. Subsequently, while most studies agree on pH being the major driver for bacterial community composition, not a single "major driver" has yet been found for fungal and protistan communities [7, 8, 45].

It logically follows that the drivers of the fungal and protistan biogeographies must be sought elsewhere. The strongest response in our models is between functional guilds - in particular, consumers strongly and consistently responding to the diversity of the preys, and more so in summer (Supplementary Table 5). To this, variation partitioning (Fig. 4) added that the combined influence of biotic and edaphic factors explained the largest share of the variance of consumer and prey communities. Thus, the intricate interplay between the environment and competitive and/or predatory interactions is better observed when functional guilds are considered.

### 4.2 Changes during snowmelt

Our results showed that alpine soil microbial communities, including protists, were progressively modified from spring to summer, with no sudden shifts during snowmelt (Supplementary Fig. 5), despite the drastic environmental changes occurring at thaw [23]. The soil microbial biomass in alpine grassland usually reaches its annual peak in winter, just before snowmelt [46]; in line with this, in our study more bacterial SSU rRNAs, which represented 98% of all sequences, were obtained from the samples collected under the snow (Fig. 3C). In soils frozen in winter, the soil microbial biomass suddenly declined at snowmelt [46, 47], and cold-adapted winter soil microbial communities died and were swiftly replaced by summer ones [48, 49]. Since we could not observe a sudden collapse of the microbial biomass at snowmelt [23], we consider that under a previous winter snow cover with stable soil temperatures above-zero (Supplementary Fig. 1), a specific winter-adapted microbial community does not develop, and thus there are no sudden biotic changes at thaw. Accordingly, we only observed changes in community composition from spring to summer.

### 4.3 Taxonomic diversity

Our study challenges two widely accepted assumptions about the diversity and the functional groups of the soil microbiome. In assessing the diversity of protists, the biases induced during PCR by so-called "universal eukaryotic primers" have already been shown: most of the PCR-primers based on the V4 region of the 18S rRNA gene overestimate taxa from the "SAR" (or Harosa) clade, the ciliates in particular, and strongly underestimate Amoebozoa (and in particular the class Conosa and the slime-moulds), as was already signalled [50, 51]. RNA-based, non-PCR, studies unequivocally agree on revealing Amoebozoa as an important (when not dominant) protistan lineage, along with Rhizaria, in soil and litter [52-54]. In our study, amoebozoans not only dominate the protistan assemblage, but also played a major role as consumers (Fig. 6) - it follows that keeping overlooking them will give only a partial view of the soil food web.

Our results indubitably showed that the bacterial predators (Myxococcota essentially) outnumbered the protistan predators. They play an essential, and mostly unrecognized, role in shaping the microbial communities (Fig. 6). Unknown biases in evaluating the rRNAs abundances cannot be ruled out - perhaps a higher abundance per cell in prokaryotes related to eukaryotes, thus overestimating the prokaryotic and underestimating the eukaryotic abundances [54] (Supplementary Note 1). Despite this, and in accordance with a previous study [55], the Myxococcota SSU rRNA dominated the predator assemblages by two orders of magnitude.

### 4.4 Methodological discussion

It increasingly appears that metatranscriptomics might become the standard for molecular monitoring of complex environmental microbiomes, for three main reasons: DNA-based surveys also include a fair portion of dead organisms: Extracellular DNA inflated the observed prokaryotic and fungal richness by up to 55% [56], and DNA-based surveys are thus inappropriate for monitoring short- to middle-term shifts. Using mock communities, it has been shown that rRNA-based methods outperform metagenomics in identifying organisms [57]. However, there are some challenges in interpreting abundances, especially because it is currently unclear how to relate rRNA transcript numbers with microbial cell numbers (Supplementary Note 1 and [50, 55, 58]).

### Conclusion

The soil microbial food web in alpine environments shows striking seasonal dynamics, with biotic interactions explaining far more variation for microbial community turnover than soil properties (e.g. pH and carbon content) or topography (elevation, spatial difference between mountains). Our study stands out by applying metatranscriptomics to a large-scale ecological assessment of the entire soil microbiota. We reached a sequencing depth overcoming the descriptive limits of classical amplicon-based approaches and making inter-domain and inter-sample comparisons possible. We have complemented our protocol with trait-based approaches to enhance the basic knowledge of the soil food web functioning. Thus, our study contributes to the understanding of the alpine ecosystem by showing the importance of the biotic interactions in shaping the seasonal changes of the soil microbiome.

## Supporting information

TableS5SummaryRDA

TableS1DatabaseEnvironmental

TableS2DatabaseTaxaTraits

TableS3TresholdSummary

TableS4SequencesObtained

## 6 Inventory of supporting information

**Supplementary Note 1**. Additional information on taxonomic diversity and methodological discussion.

**Supplementary Table 1**. Database of the topographic, edaphic and biotic parameters of the samples.

**Supplementary Table 2**. Database of the abundance of each identified taxon per sample, with their taxonomic and functional assignment.

**Supplementary Table 3**. Thresholds applied to select the most abundant taxa at different taxonomic levels, and the number of taxa and percentage of sequences deleted.

**Supplementary Table 4**. Sequences obtained from the 158 soil samples and results at each step of their processing.

**Supplementary Table 5**. Models of the most influential topographic, edaphic and biotic variables, selected by distance-based redundancy and variable selection (functions dbrda and ordistep), for consumers and preys and also bacteria, protists and fungi. Models evaluated by anova, F-values and model p-values are shown.

**Supplementary Fig. 1**. Soil temperature recorded every two hours from autumn 2019 to summer 2020, averaged by altitude categories. In winter the soil remained at a constant temperature just above zero degrees, except in the highest sites, where temperatures dropped slightly below 0°C but stayed above -1°C.

**Supplementary Fig. 2**. Principal Component Analysis of the Bray-Curtis dissimilarity indices of the main taxa and functions, showing that snow coverage, season, altitude and mountain have little influence in shaping the communities. The functional group "preys", was nearly identical to bacteria, and thus not shown.

**Supplementary Fig. 3**. Rarefaction curves for bacteria, protists and fungi, by snow cover, altitude and mountain. They were calculated using the iNEXT package, on raw abundances, with a 97% confidence interval, 50 bootstraps and 50 knots.

**Supplementary Fig. 4**. Accumulation curves for bacteria, protists and fungi, by snow cover, altitude and mountain. They were calculated using the *specaccum* function, on raw abundances.

**Supplementary Fig. 5**. Variation in the mean of SSU sequences, diversity (Shannon index) and evenness, from the samples under the snow, at the edge of the snow patch and snow-free. There were no significant changes (Tukey-test, p-value ≤ 0.05), they are indicated by "a". Standard errors bars are shown. **a**, Consumers (predatory bacteria, heterotrophic and free-living protists, selected nematodes). **b**, Preys (non-predatory bacteria, fungi and autotrophic protists).

**Supplementary Fig. 6**. Co-occurrence networks of abundant phyla of preys. **a**, Spring, under the snow. **b,** Summer. The size of the nodes (dots) are proportional to the number reads. Edges (connecting lines) represent positive (light grey) or negative (dark grey) correlations, with line width proportional to the number of correlations. Self-loops and taxa with a single edge are not shown.

## 7 Acknowledgments

For help in collecting samples and doing the vegetation survey in Davos, we are grateful to Sven Buchmann, Fiona Schwaller, Sabrina Keller and Francesca Jaroszynska. We thank Jana Täumer and Verena Gross at the laboratory at the University of Greifswald, for helping with RNA extraction. At the University of Cologne, we are grateful to Irene Brockhaus, Anna Herzog, Andrea Meis, and Oscar Rindt for the measurement of edaphic properties, under the supervision of Christoph Rosinger. We are indebted to Caterina Penone for conducting the variance partitioning analysis. We thank Alexandra Elbakyan for facilitating access to scientific articles.

## 8 Author contribution

AMFD and CR collected samples. CR conducted the vegetation survey. AMFD and TU performed molecular biological analyses. AMFD and MBD conducted bioinformatic analyses. AMFD, JF, MB and TU and performed statistical analyses. AMFD wrote the manuscript. All authors contributed to the interpretation of the data and the writing/editing of the manuscript. AMFD devised and coordinated the project.

## 9 Funding

This work was funded by the German Research Foundation (DFG), Grant Number FI 1686/2-1. Open access funding enabled and organized by Projekt DEAL.

## 10 Data Availability

Sequencing data and raw sequences are available under NCBI BioProject PRJNA850398.

## 11 Declarations

*Ethics approval and consent to participate*

Not applicable

*Consent for publication*

Not applicable

*Competing interests*

The authors declare that they have no competing interests.

## Notes

### Competing Interest Statement

The authors have declared no competing interest.

## References

1. Cheng L, Zhang N, Yuan M, Xiao J, Qin Y, Deng Y, Tu Q, Xue K, Van Nostrand JD, Wu L et al: Warming enhances old organic carbon decomposition through altering functional microbial communities. ISME J 2017, 11:1825–1835.

2. Zhu B, Li C, Wang J, Li J, Li X: Elevation rather than season determines the assembly and co-occurrence patterns of soil bacterial communities in forest ecosystems of Mount Gongga. Appl Microbiol Biotechnol 2020, 104:7589–7602.

3. Shen C, He J-Z, Ge Y: Seasonal dynamics of soil microbial diversity and functions along elevations across the treeline. Sci Total Environ 2021, 794:148644.

4. Adamczyk M, Hagedorn F, Wipf S, Donhauser J, Vittoz P, Rixen C, Frossard A, Theurillat J-P, Frey B: The soil microbiome of GLORIA mountain summits in the Swiss Alps. Front Microbio 2019, 10:1080.

5. Fiore-Donno AM, Richter-Heitmann T, Degrune F, Dumack K, Regan KM, Marhan S, Boeddinghaus RS, Rillig M, Friedrich MW, Kandeler E et al: Functional traits and spatio-temporal structure of a major group of soil protists (Rhizaria: Cercozoa) in a temperate grassland. Front Microbio 2019, 10:1332.

6. Yashiro E, Pinto-Figueroa E, Buri A, Spangenberg JE, Adatte T, Niculita-Hirzel H, Guisan A, van der Meer JR: Local environmental factors drive divergent grassland soil bacterial communities in the Western Swiss Alps. Appl Environ Microbiol 2016, 82:6303–6316.

7. Shen C, Gunina A, Luo Y, Wang J, He J-Z, Kuzyakov Y, Hemp A, Classen AT, Ge Y: Contrasting patterns and drivers of soil bacterial and fungal diversity across a mountain gradient. Environ Microbiol Rep 2020, 22:3287–3301.

8. Mazel F, Malard L, Niculita-Hirzel H, Yashiro E, Mod HK, Mitchell EAD, Singer D, Buri A, Pinto E, Guex N et al: Soil protist function varies with elevation in the Swiss Alps. Environ Microbiol Rep 2021, 24:1689–1702.

9. Nguyen B-AT, Chen Q-L, Yan Z-Z, Li C, He J-Z, Hu H-W: Distinct factors drive the diversity and composition of protistan consumers and phototrophs in natural soil ecosystems. Soil Biol Biochem 2021, 160:108317.

10. Li G, Wang Y, Lib H, Zhang X, Gong J: Quantifying relative contributions of biotic interactions to bacterial diversity and community assembly by using community characteristics of microbial eukaryotes. Ecological Indicators 2023, 146:109841.

11. Lee CK, Laughlin DC, Bottos EM, Caruso T, Joy K, Barrett JE, Brabyn L, Nielsen UN, Adams BJ, Wall DH et al: Biotic interactions are an unexpected yet critical control on the complexity of an abiotically driven polar ecosystem. Comm biol 2019, 2:62.

12. García-Girón J, Heino J, García-Criado F, Fernández-Aláez C, Alahuhta J: Biotic interactions hold the key to understanding metacommunity organisation. Ecography 2020, 43:1180–1190.

13. Chesson P, Kuang JJ: The interaction between predation and competition. Nature 2008, 456:235–238.

14. Dormann CF, Bobrowski M, Dehling DM, Harris DJ, Hartig F, Lischke H, Moretti MD, Page J, Pinkert S, Schleuning M et al: Biotic interactions in species distribution modelling: 10 questions to guide interpretation and avoid false conclusions. Global Ecol Biogeogr 2018, 27(9):1004–1016.

15. Federal Office for Meteorology and Climatology. [www.meteoswiss.admin.ch/home/climate/swiss-climate-in-detail.html]. Last accessed March 2023.

16. Swiss Confederation, Geocatalog. [https://map.geo.admin.ch/]. Last accessed March 2023.

17. Gruber-Vodicka HR, Seah BK, Pruesse E: phyloFlash: Rapid small-subunit rRNA profiling and targeted assembly from metagenomes mSystems 2020, 5:e00920–00920.

18. Magoč T, Salzberg SL: FLASH: fast length adjustment of short reads to improve genome assemblies. Bioinformatics 2011, 27:2957–2963.

19. Guillou L, Bachar D, Audic S, Bass D, Berney C, Bittner L, Boutte C, Burgaud G, de Vargas C, Decelle J et al: The Protist Ribosomal Reference database (PR2): a catalog of unicellular eukaryote small subunit rRNA sequences with curated taxonomy. Nucleic Acids Res 2013, 41:D597–D604.

20. Camacho C, Coulouris G, Avagyan V, Ma N, Papadopoulos J, Bealer K, Madden TL: BLAST+: architecture and applications. BMC Bioinf 2008, 10:421.

21. Yeates G, Bongers T, De Goede R, Freckman D, Georgieva S: Feeding habits in soil nematode families and genera - an outline for soil ecologists. J Nematol 1993, 25:315–331.

22. Nguyen NH, Song Z, Bates ST, Branco S, Tedersoo L, Menke J, Schilling JS, Kennedy PG: FUNGuild: An open annotation tool for parsing fungal community datasets by ecological guild. Fungal Ecol 2016, 20:241–248.

23. Rindt O, Rosinger C, Bonkowski M, Rixen C, Brüggemann N, Urich T, Fiore-Donno AM: Biogeochemical dynamics during snowmelt and in summer in the Alps. Biogeochemistry 2023, 162:257–266.

24. Juillerat P, Bäumler B, Bornand C, Gygax A, Jutzi M, Möhl A, Nyffeler R, Sager L, Santiago H, Eggenberg S: Flora Helvetica Checklist 2017: der Gefässpflanzen der Schweiz. Geneva: InfoFlora; 2017.

25. R Development Core Team: R: A language and environment for statistical computing. In. Edited by Computing RFfS. Vienna, Austria; 2014.

26. Oksanen J, Blanchet FG, Kindt R, Legendre P, Minchin PR, O’Hara RB, Simpson GL, Solymos P, Stevens MHH, Wagner H: Vegan: Community Ecology Package. R package version 2.0-10. http://CRAN.R-project.org/package=vegan. 2013.

27. Love MI, Huber W, Anders S: Moderated estimation of fold change and dispersion for RNA-seq data with DESeq2. Genome Biol 2014, 15:550.

28. Tackmann J, Rodrigues JFM, von Mering C: Rapid inference of direct interactions in large-scale ecological networks from heterogeneous microbial sequencing data. Cell Systems 2019, 9:286–296.

29. Bezanson J, Karpinski S, Shah VB, Edelman A: Julia: A fast dynamic language for technical computing. arXiv 2012, 1209:5145.

30. Shannon P, Markiel A, Ozier O, Baliga N, Wang J, Ramage D, Amin N, Schwikowski B, Ideker T: Cytoscape: a software environment for integrated models of biomolecular interaction networks. Genome Res 2003, 13:2498–2504.

31. Gauzens B, Legendre S, Lazzaro X, Lacroix G: Intermediate predation pressure leads to maximal complexity in food webs. Oikos 2015, 125:595–603.

32. Saleem M, Fetzer I, Dormann CF, Harms H, Chatzinotas A: Predator richness increases the effect of prey diversity on prey yield. Nat Commun 2012, 3:1305.

33. Burian A, Pinn D, Peralta-Maraver I, Sweet M, Mauvisseau Q, Eyice O, Bulling M, Röthig T, Kratina P: Predation increases multiple components of microbial diversity in activated sludge communities. ISME J 2022, 16:1086–1094.

34. Corno G, Caravati E, Callieri C, Bertoni R: Effects of predation pressure on bacterial abundance, diversity, and size-structure distribution in an oligotrophic system. J Limnol 2008, 67:107–119.

35. Wittebolle L, Marzorati M, Clement L, Balloi A, Daffonchio D, Heylen K, De Vos P, Verstraete W, Boon N: Initial community evenness favours functionality under selective stress. Nature 2009, 458:623–626.

36. Crowder DW, Northfield Td, Strand MR, Snyder WE: Organic agriculture promotes evenness and natural pest control. Nature 2010, 466:109–113.

37. Tveit A, Urich T, Svenning MM: Metatranscriptomic analysis of Arctic peat soil microbiota. Appl Environ Microbiol 2014, 80:5761.

38. Rossmann M, Pérez-Jaramillo J, Kavamura V, Chiaramonte J, Dumack K, Fiore-Donno AM, Mendes LW, Ferreira M, Bonkowski M, Raaijmakers JM et al: Multitrophic interactions in the rhizosphere microbiome of wheat: from bacteria and fungi to protists. FEMS Microbiol Ecol 2020, 96:fiaa032.

39. Rüger L, Feng K, Dumack K, Freudenthal J, Chen Y, Sun R, Wilson M, Yu P, Sun B, Deng Y et al: Assembly patterns of the rhizosphere microbiome along the longitudinal root axis of maize (Zea mays L.). Front Microbio 2021, 12:614501.

40. Fiore-Donno AM, Human ZR, Štursová M, Mundra S, Morgado LN, Kauserud H, Baldrian P, Bonkowski M: Soil compartments (bulk soil, litter, root and rhizosphere) as main drivers of soil protistan communities distribution in forests with different nitrogen deposition. Soil Biol Biochem 2022, 168:108628.

41. Schostag M, Priemé A, Jacquiod S, Russel J, Ekelund F, Jacobsen CS: Bacterial and protozoan dynamics upon thawing and freezing of an active layer permafrost soil. ISME J 2019, 13:1345–1359.

42. Eckert EM, Baumgartner M, Huber IM, Pernthaler J: Grazing resistant freshwater bacteria profit from chitin and cell-wall-derived organic carbon. Environ Microbiol Rep 2013, 15:2019–2030.

43. Romdhane S, Spor A, Aubert J, Bru D, Breuil M-C, Hallin S, Mounier A, Ouadah S, Tsiknia M, Philippot L: Unraveling negative biotic interactions determining soil microbial community assembly and functioning. ISME J 2022, 16:296–306.

44. Zheng J, Dini-Andreote F, Luan L, Geisen S, Xue J, Li H, Sun B, Jiang Y: Nematode predation and competitive interactions affect microbe-mediated phosphorus dynamics. MBio 2022, 13(3):1–13.

45. Li J, Li C, Kou Y, Yao M, He Z, Li X: Distinct mechanisms shape soil bacterial and fungal co-occurrence networks in a mountain ecosystem. FEMS Microbiol Ecol 2020, 96:fiaa030.

46. Schmidt SK, Lipson DA: Microbial growth under the snow: Implications for nutrient and allelochemical availability in temperate soils. Plant Soil 2004, 259:1–7.

47. Ernakovich JG, Hopping KA, Berdanier AB, Simpson RT, Kachergis EJ, Steltzer H, Wallenstein MD: Predicted responses of arctic and alpine ecosystems to altered seasonality under climate change. Glob Change Biol 2014, 20:3256–3269.

48. Buckeridge KM, Banerjee S, Siciliano SD, Grogan P: The seasonal pattern of soil microbial community structure in mesic low arctic tundra. Soil Biol Biochem 2013, 65:338–347.

49. Broadbent AA, Snell HS, Michas A, Pritchard WJ, Newbold L, Cordero I, Goodall T, Schallhart N, Kaufmann R, Griffiths RI et al: Climate change alters temporal dynamics of alpine soil microbial functioning and biogeochemical cycling via earlier snowmelt. ISME J 2021, 15:2264–2275.

50. Geisen S, Tveit A, Clark IM, Richter A, Svenning MM, Bonkowski M, Urich T: Metatranscriptomic census of active protists in soils. ISME J 2015, 9: 2178–2190.

51. Fiore-Donno AM, Weinert J, Wubet T, Bonkowski M: Metacommunity analysis of amoeboid protists in grassland soils. Sci Rep 2016, 6:19068.

52. Urich T, Lanzén A, Qi J, Huson DH, Schleper C, Schuster SC: Simultaneous assessment of soil microbial community structure and function through analysis of the meta-transcriptome. PLoS ONE 2008, 3(6):e2527.

53. Voss C, Fiore-Donno AM, Guerreiro MA, Peršoh D, Bonkowski M: Metatranscriptomics reveal unsuspected protistan diversity in leaf litter across temperate beech forests, with Amoebozoa the dominating lineage. FEMS Microbiol Ecol 2019, 95:fiz142.

54. Harkes P, Suleiman AKA, van den Elsen SJJ, de Haan JJ, Holterman M, Kuramae EE, Helder J: Conventional and organic soil management as divergent drivers of resident and active fractions of major soil food web constituents. Sci Rep 2019, 9:13521.

55. Petters S, Groß V, Söllinger A, Pichler M, Reinhard A, Bengtsson MM, Urich T: The soil microbial food web revisited: Predatory myxobacteria as keystone taxa? ISME J 2021, 15:2665–2675.

56. Carini P, Marsden PJ, Leff J, Morgan EE, Strickland MS, Fierer N: Relic DNA is abundant in soil and obscures estimates of soil microbial diversity. Nat Microbiol 2016, 2:16242.

57. Hempel CA, Wright N, Harvie J, Hleap JS, Adamowicz SJ, Steinke D: Metagenomics versus total RNA sequencing: most accurate data-processing tools, microbial identification accuracy and perspectives for ecological assessments. Nucleic Acids Res 2022, 50:9279–9293.

58. Söllinger A, Séneca J, Borg Dahl M, Motleleng LL, Prommer J, Verbruggen E, Sigurdsson BD, Janssens I, Peñuelas J, Urich T et al: Down-regulation of the bacterial protein biosynthesis machinery in response to weeks, years, and decades of soil warming. Sci Adv 2022, 8:eabm3230.

