## SupplementaryFigures_1_6 for "Biotic interactions explain seasonal dynamics of the alpine soil microbiome"

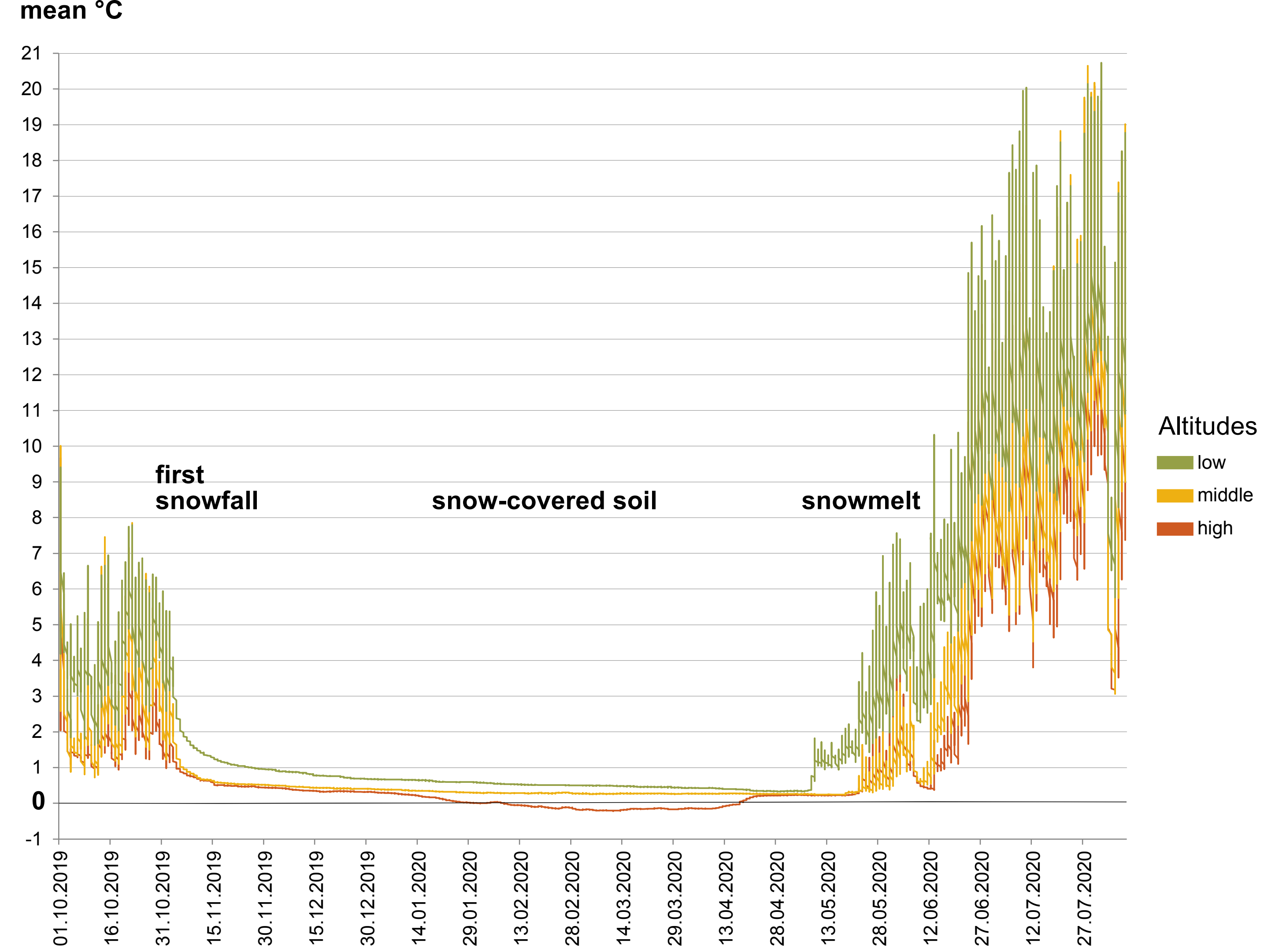


**Figure S1.** Soil temperature recorded every two hours from autumn 2019 to summer 2020, averaged by altitude categories. In winter the soil remained at a constant temperature just above zero degrees, except in the highest sites, where temperatures dropped slightly below 0°C but stayed above -1°C.

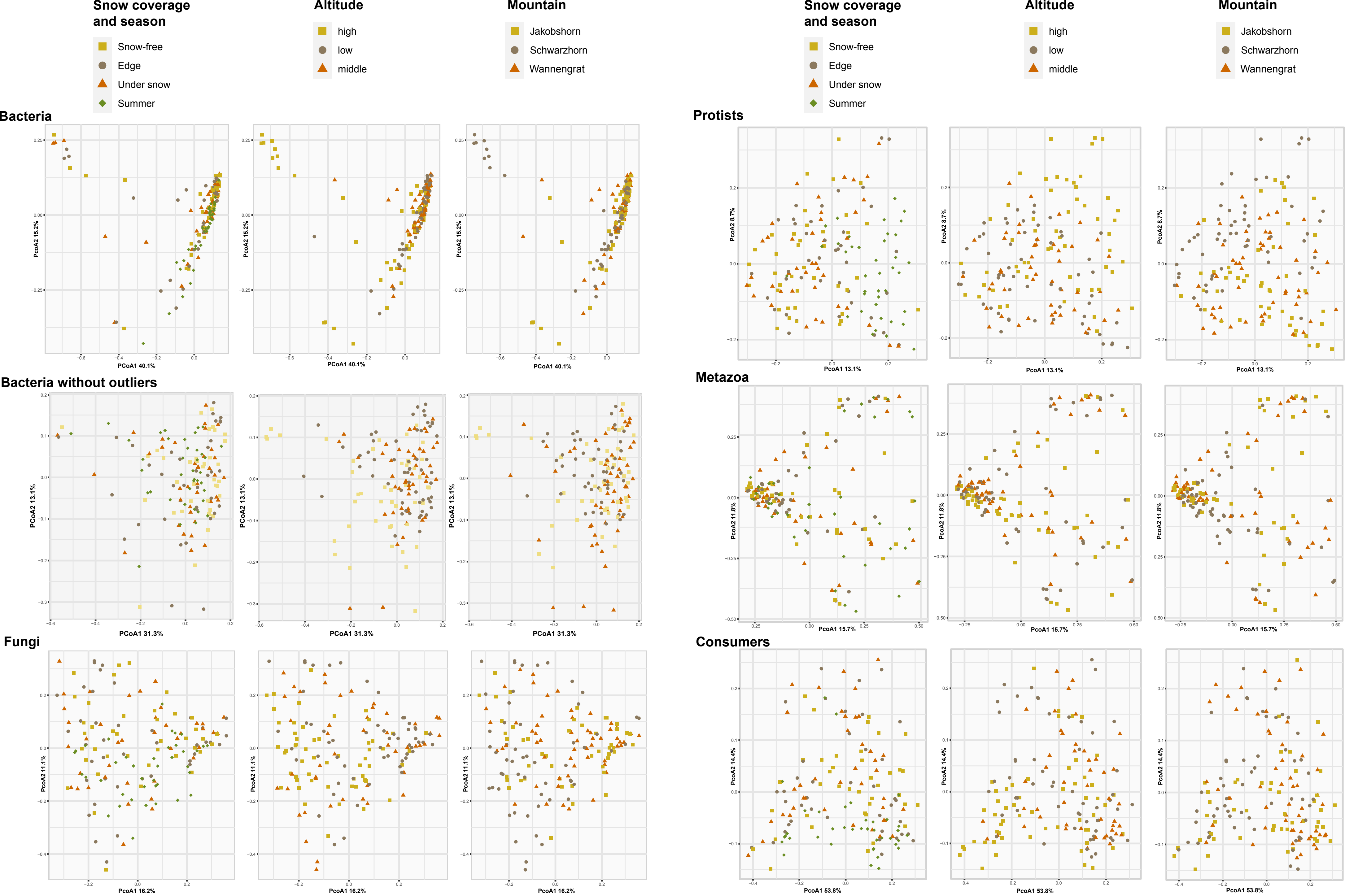


**Figure S2**. Principal Component Analysis of the Bray-Curtis dissimilarity indices of the main taxa and functions, showing that snow coverage, season, altitude and mountain have little influence in shaping the communities. The functional group "preys" was nearly identical to bacteria, and thus not shown.

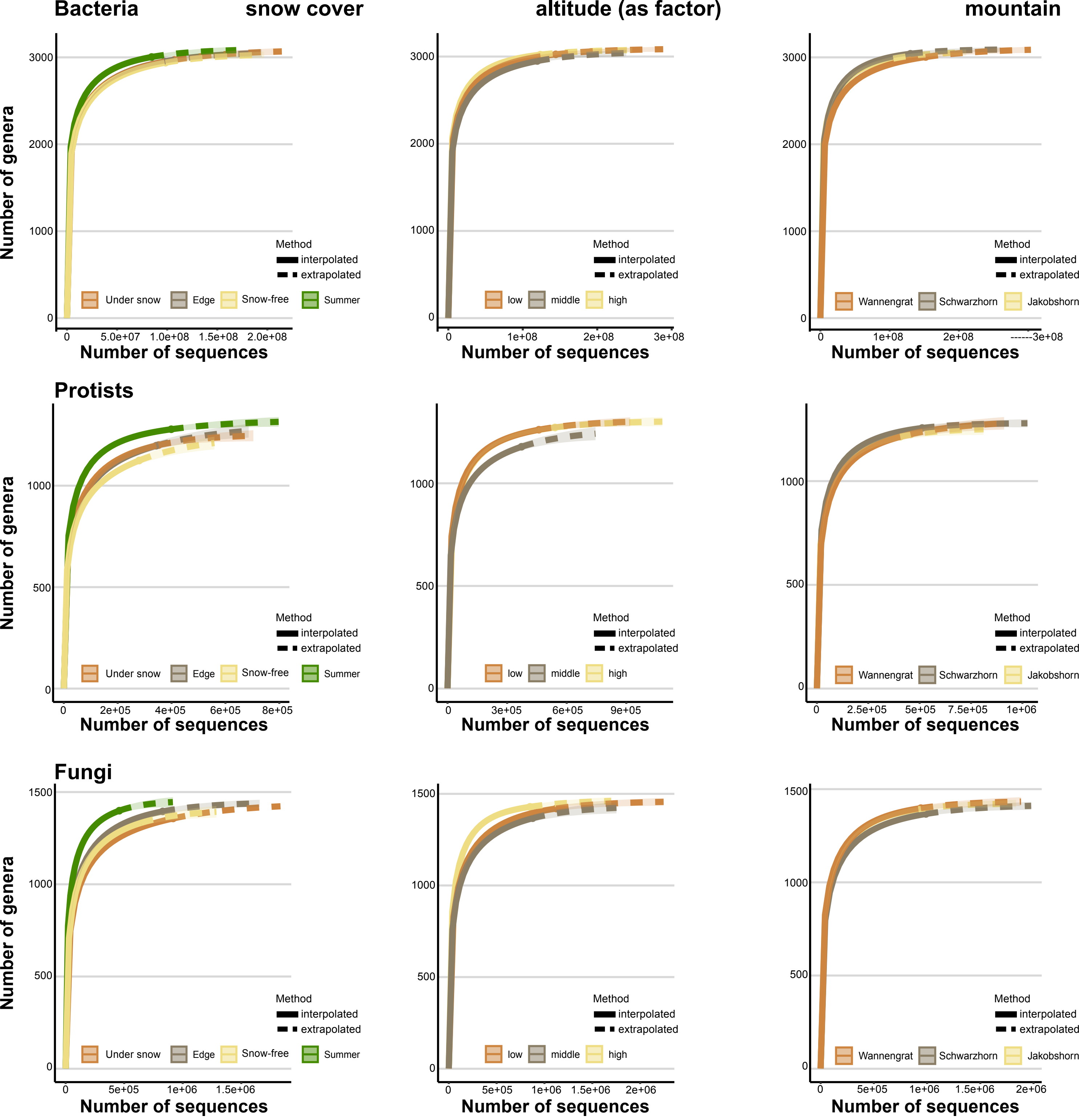


**Figure S3**. Rarefaction curves for bacteria, protists and fungi, by snow cover, altitude and mountain. They were calculated using the iNEXT package, on raw abundances, with a 97% confidence interval, 50 bootstraps and 50 knots.

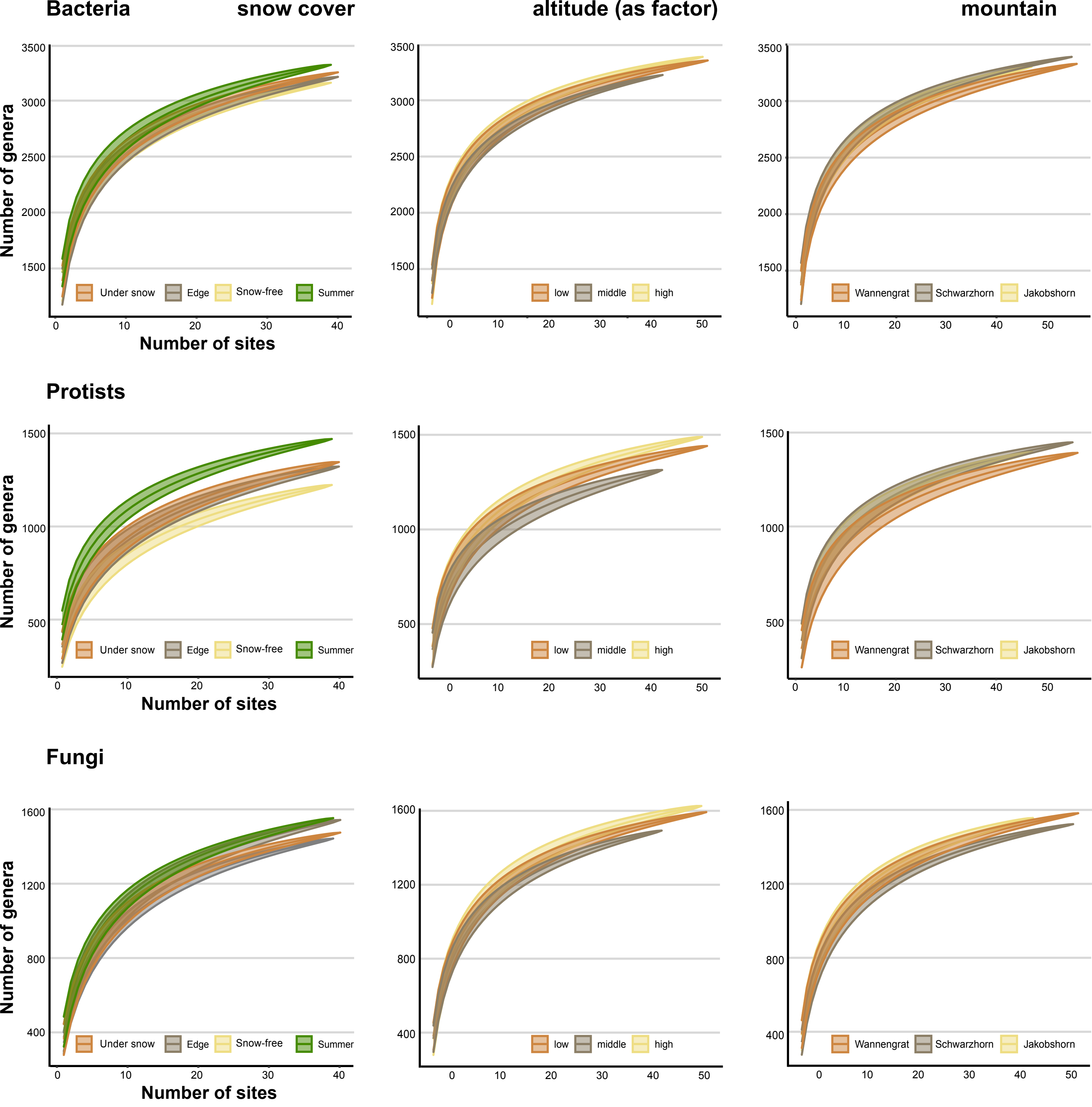


**Figure S4**. Accumulation curves for bacteria, protists and fungi, by snow cover, altitude and mountain. They were calculated usin the *specaccum* function, on raw abundances.

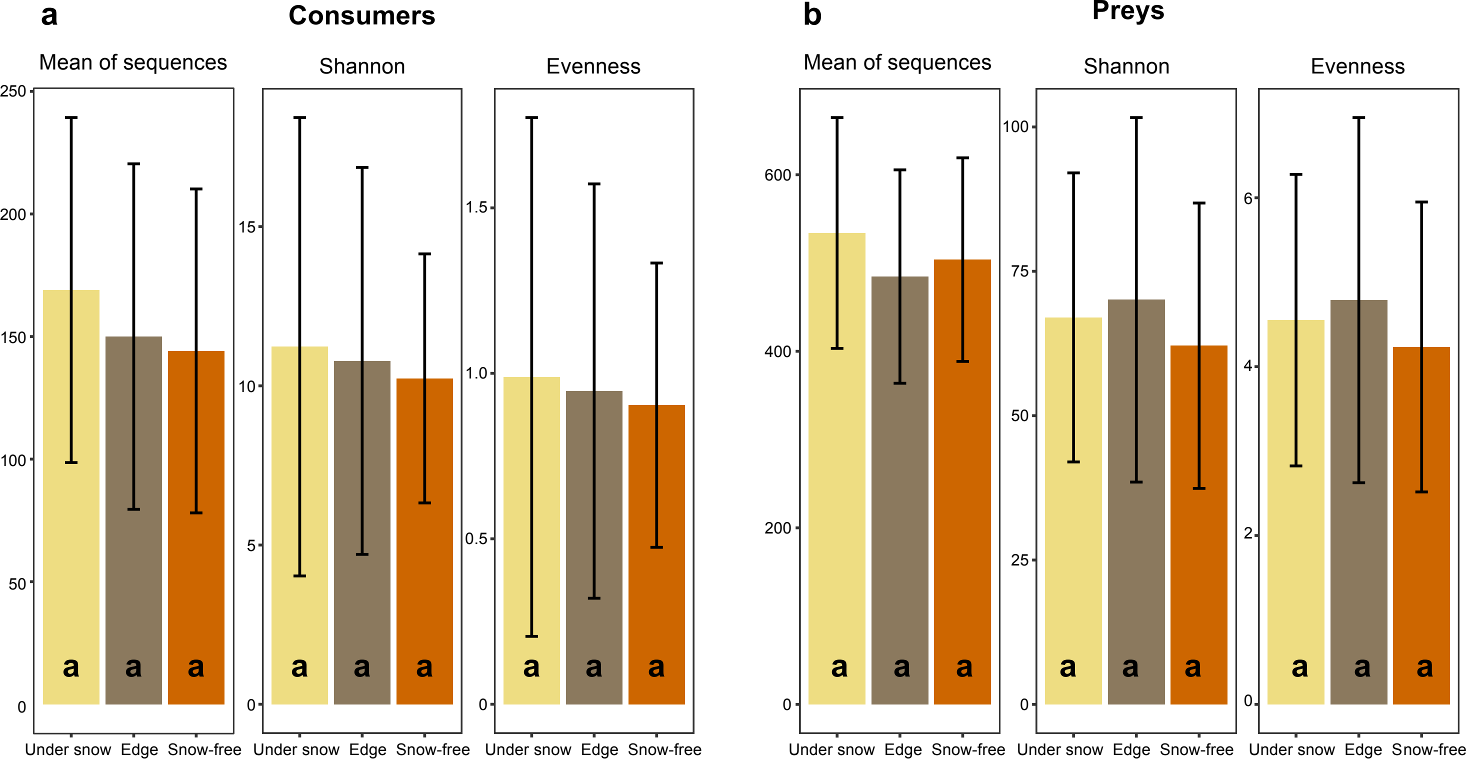


**Figure S5**. Variation in the mean of SSU sequences, diversity (Shannon index) and evenness, from the samples under the snow, at the edge of the snow patch and snow-free. There were no significant changes (Tukey-test, p-value⩽0.05), they are indicated by "a". Standard errors bars are shown. **a**, Consumers (predatory bacteria, heterotrophic and free-living protists, selected nematoda). **b**, Preys (non-predatory bacteria, fungi and autotrophic protists).

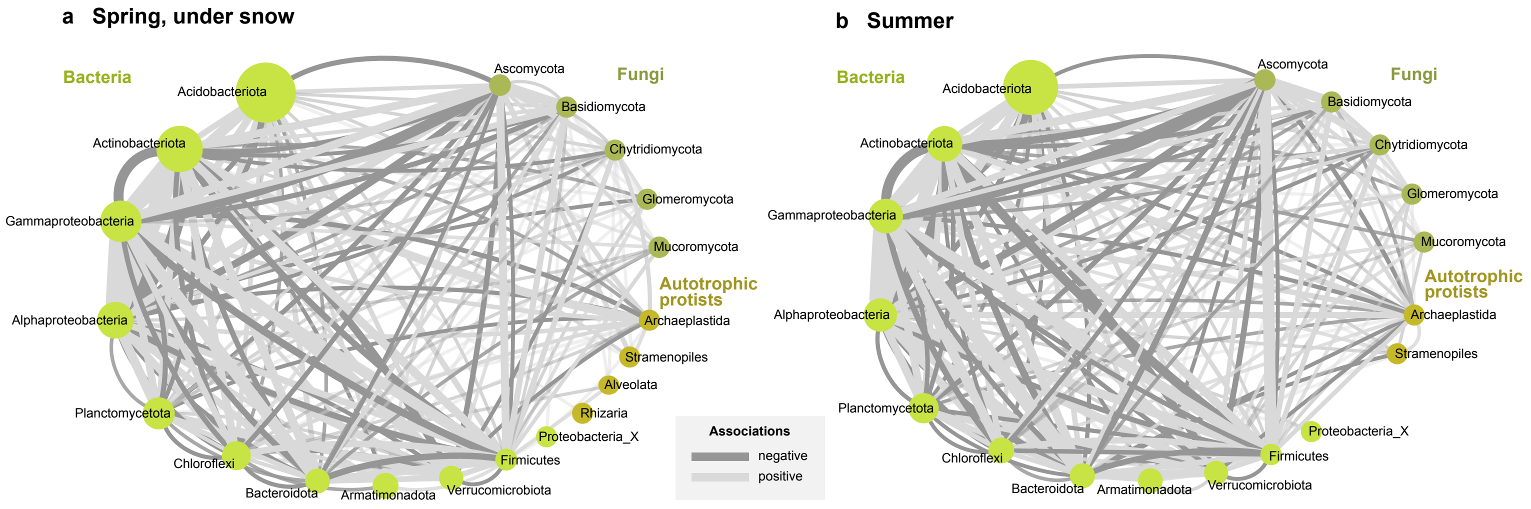


**Figure S6**. Co-occurrence networks of abundant phyla of preys. **a,** Spring, under the snow. **b**, Summer. The size of the nodes (dots) are proportional to the number reads. Edges (connecting lines) represent positive (light grey) or negative (dark grey) correlations, with line width proportional to the number of correlations. Self-loops and taxa with a single edge are not shown.
